# Single-cell spatial multi-omic characterization of the tumour microenvironment in transformed follicular lymphoma

**DOI:** 10.64898/2026.08.11.743383

**Authors:** Shaocheng Wu, Eric Lee, Anne-Sophie Fratzscher, Andrew Lytle, Spencer D. Martin, Tina Hsu, Adele Telenius, Yifan Yin, Amos Fong, Shinya Rai, Manuba Fujisawa, Honor Cheung, Samuel Aparicio, David G. Huntsman, Tomohiro Aoki, David W. Scott, Christian Steidl, Andrew Roth

## Abstract

Histological examination of follicular lymphoma (FL) biopsies remains the cornerstone for diagnostic grading of FL. Single-cell sequencing approaches, while transcriptomically rich, require tissue dissociation and lose the native spatial context that underpins FL transformation to diffuse large B-cell lymphoma (DLBCL). To investigate the spatial interplay between malignant B-cells and the tumour microenvironment (TME) across disease states, we performed subcellular single-cell spatial transcriptomics and spatial proteomics on 12 paired pre/post-transformation samples and 10 non-transforming FL controls, integrated with matched single-cell whole genome sequencing (scWGS). Our analysis reveals that transformation is accompanied by a shift toward B-cell-predominant stromal and immunosuppressive cellular neighbourhoods, where the magnitude of expansion correlates with time to transformation. Prior to transformation, immunomodulatory Galectin-9 interactions move from the intra-follicular core to the extra-follicular space. Integration with scWGS demonstrates that high copy-number instability in malignant B-cells is associated with reduced supportive T-cell niches and intensified immunoregulatory crosstalk at the transformed state. Collectively, our multi-omic analysis characterizes TME remodeling during FL transformation, contributing to a refined disease evolution model.

## Introduction

Follicular lymphoma (FL) is the second most common non-Hodgkin lymphoma and the most frequent indolent type. While typically considered incurable, it is characterized by a favorable overall survival often surpassing 10 years^1–3^. However, FL exhibits a highly variable clinical course where most patients eventually progress, and a subset of patients experiences histological transformation to an aggressive lymphoma, frequently diffuse large B-cell lymphoma (DLBCL). Histological transformation of FL is observed in 2 to 3 percent of patients per year^4^, represented by a diffuse proliferation of neoplastic centroblasts that efface the follicular structures reminiscent of germinal centres^5^. The disruption of the tumour microenvironment (TME) architecture during transformation involves a malignant B-cell facilitated reprogramming of the immunophenotype to feature a pro-tumour microenvironment^6,7^. Transformation is associated with significantly worse prognosis, marked by rapid disease progression and reduced overall survival^8^. Despite this, no definitive clinical biomarkers have been established to reliably predict transformation, reflecting the inherent challenges in deconvoluting the phenotypic complexity of the TME and its interactions with malignant components.

Previous studies using flow cytometry and immunohistochemistry (IHC) have demonstrated that specific TME cell populations, including cytotoxic T cells and macrophages, are associated with delayed disease progression and improved survival outcomes^9,10^. More recently, single-cell techniques have been leveraged to probe the genotypic landscape, malignant cell phenotypic evolution and TME heterogeneity in FL^11,12^. Temporal analysis on paired pre and post-transformed FL samples has further revealed a shift towards an exhausted and expansively regulatory immunophenotype during transformation^13^. This is exemplified by malignant cells often inducing an increase in regulatory T cell subsets to inhibit cytotoxic activity of effector cells^14^. Additionally, specific lymph node stromal cell populations have been described to upregulate genes involved in extracellular matrix (ECM) remodeling and co-localize with malignant cells to assist in avoiding immune surveillance^15^. While the cellular prevalence of various TME cell states offers potential prognostic value^16^, their spatial interplay within the FL TME architecture remains to be fully characterized, highlighting an ideal opportunity for the use of emerging high-throughput spatial transcriptomic methods.

In this study, we leverage single-cell spatial transcriptomics to map the landscape of TME dynamics across various disease states, including FL from patients who have not experienced transformation (non-tFL) and paired pre-transformed FL (tFL-FL) and post-transformed FL (tFL-DLBCL) samples. We additionally integrated orthogonal single-cell spatial proteomics for protein-level validation alongside matched single-cell DNA and RNA sequencing to comprehensively annotate and contrast spatial features associated with transformation. Our analysis reveals that FL transformation exhibits a reorganization of cellular neighbourhoods and intercellular communication networks in the TME. The architectural transition is highlighted by a spatial shift of Galectin-9–TIM-3 interactions from the intra-follicular zone to the extra-follicular space, and the emergence of genomic instability in the malignant-B cells as a feature of the transformed TME closely linked to disrupted B–T-CM (central memory T cell) neighbourhoods in an immunoregulatory microenvironment.

## Results

### 1 Spatially resolved single-cell profiling of the transformed FL cellular ecosystem

To profile and investigate the spatial cellular heterogeneity of FL transformation, we generated a single-cell spatial dataset comprising 12 FL patients with matched excisional biopsies at pre-and post-transformation disease states (pre: tFL-FL, post: tFL-DLBCL) and 10 FL patients with no disease progression or transformation within six years (non-tFL), and one additional transformed patient with a post-transformation (tFL-DLBCL) biopsy only (tFL10) [Supplementary Fig.1a,b]. An overview of the study is shown in [Fig.1a], and clinical details of the cohort are provided in Supplementary Table 1. A tissue microarray (TMA) constructed of the 23 patients was subjected to in-situ subcellular multiplexed transcriptome profiling using the CosMx Spatial Molecular Imager (SMI) platform. The Human 6K Discovery Panel with 200 add-on genes was used for imaging, incorporating B-cell lymphoma-specific genes to capture cell states and cellular interactions (Supplementary Table 2).

**Fig 1.**
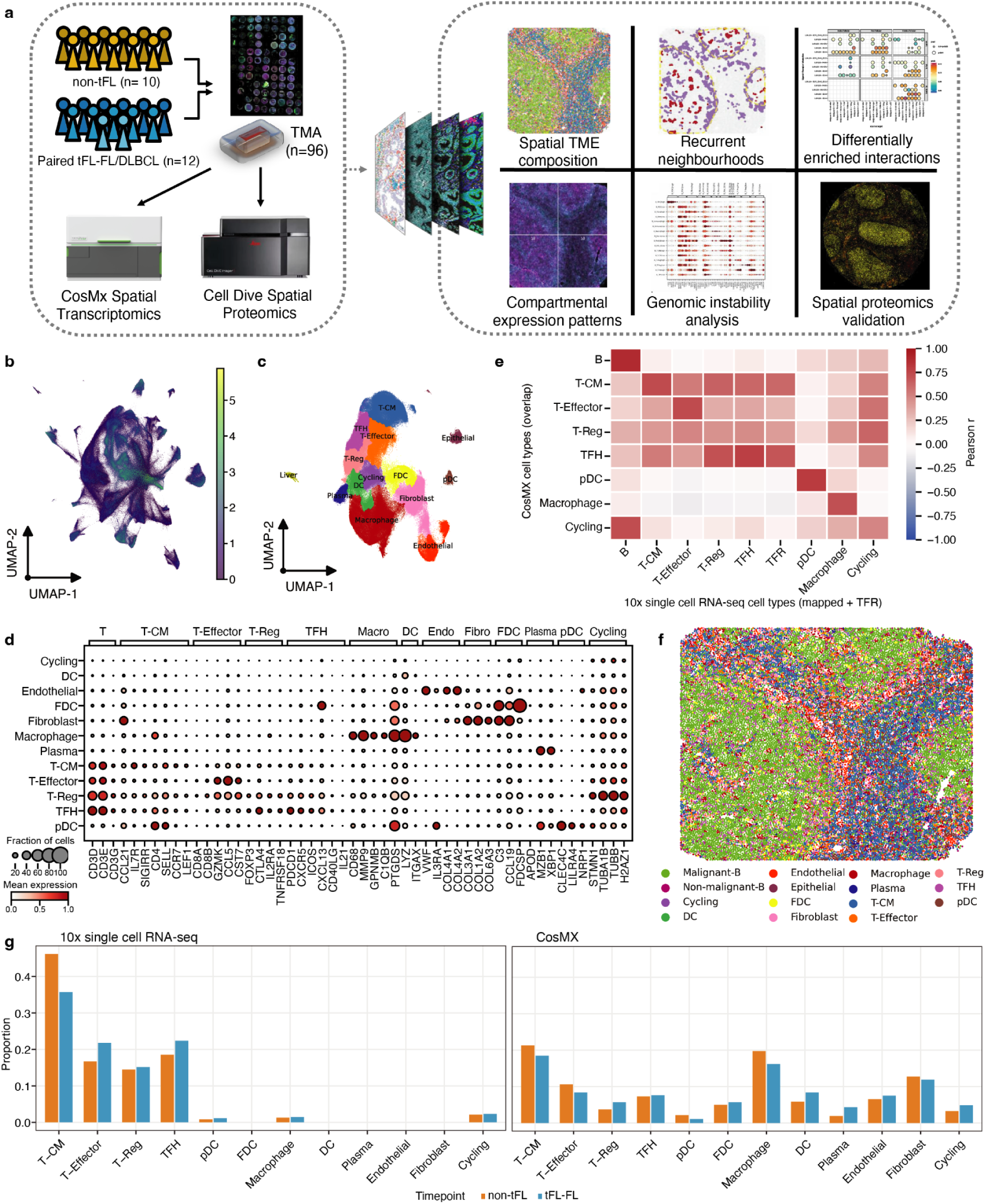
Overview of spatially resolved single-cell profiling of the transformed FL cellular ecosystem. **a,** Overview of high-dimensional spatial imaging of the FL cellular ecosystem and the analysis of the cellular composition in the FL TME between disease states. **b,** UMAP of all cells colored by the expression of MS4A1. **c,** UMAP of all cells colored by cell type. **d,** Dotplot representing the mean expression of transcripts in cell types. **e,** Heatmap of the correlation between marker expression of cell types identified in the 10x scRNA-seq dataset and the CosMx dataset. **f,** Spatial distribution of cell type annotations in a representative core of a non-tFL patient. **g,** Barplot indicating the proportion of cell types across all cells in the 10x scRNA-seq dataset and the CosMx dataset.

Transcripts were decoded using vendor supplied software across over 300 fields of view (FOV) and mapped onto cell masks generated from cell segmentation. We identified a total of 959,551 cells with a median of 13,707 cells per core (non-tFL:17,168; tFL-FL:14,135; tFL-DLBCL:11,591) and a median of 471.2 genes detected per cell following quality control filtering. To provide orthogonal validation at the protein-level, we additionally acquired 21-plex single-cell proteomic measurements using the Cell DIVE multiplexed imaging solution. These data were integrated with previously collected modalities, including single-cell whole genome sequencing (scWGS) and single-cell RNA sequencing (scRNA-seq) on the same cohort. To streamline computational analysis, we applied a unified analytical framework to perform end-to-end analysis from image processing to spatial modeling^40^.

Following the cell type annotation (see methods), we identified 14 distinct cell types spanning malignant B-cells (Malignant-B, n=463,835), non-malignant B-cells (Non-malignant-B, n=88,530), macrophages (Macrophage, n=77,293), central memory T cells (T-CM, n=63,609), fibroblasts (Fibroblast, n=43,637), effector T cells (T-Effector, n=38,836), follicular T helper cells (TFH, n=28,843), regulatory T cells (T-Reg, n=25,700), endothelial cells (Endothelial, n=23,875), dendritic cells (DC, n=21,948), follicular dendritic cells (FDC, n=19,236), benign cycling cells (Cycling, n=17,416), plasma cells (Plasma, n=6,423), and plasmacytoid dendritic cells (pDC, n=5,448) [Fig.1c and 1d]^13^. The cell type annotations were validated through a series of measures. Briefly, cell types were assessed for similarity in gene signatures to matched scRNA-seq measurements from the same cohort. Gene expression profiles of commonly identified cell types across the two transcriptomic modalities were concordant (r=0.71, Pearson correlation), with a correlation magnitude comparable to recent observations between disaggregated and spatial measurements of the same tissue^17^ [Fig.1e]. Additionally, cells were spatially mapped to verify that their localization aligned with the pathobiological features observed in corresponding H&E-stained sections [Fig.1f and Supplementary Fig.1c,d]. Our analysis revealed a strong discordance in cell types captured between disaggregate and spatial modalities. The CosMx dataset revealed an overall increase in macrophages and enabled the detection of DCs, FDCs, plasma cells, endothelial cells, and fibroblasts that were not detected in our scRNA-seq dataset [Fig.1g]. The differences in cell type distribution are likely attributable to imaging-based spatial transcriptomics, such as CosMx, bypassing cell dissociation and avoids the cell size biases inherent to droplet-based scRNA-seq^18^, yielding a more accurate representation of the FL TME.

### 2 Spatial profiling reveals shifts in cellular neighbourhoods in transformed FL

Previous scRNA-seq studies have illustrated evolutionary changes in TME composition during FL transformation, particularly within T cell proportions^13^. Leveraging our spatial data, we sought to investigate whether the single-cell compositional shifts and colocalization patterns were maintained or amplified between non-tFL and tFL-FL states. We observed a higher frequency of T-Regs and lower frequency of T-CM and TFH among the T cell populations in tFL-FL compared to non-tFL, despite observing no significant differences in overall distribution of cell types (p > 0.05, Wilcoxon rank-sum test) [Fig.1g].

We next considered whether cellular neighbourhoods (CNs) differed, as cell behavior is shaped by interactions within local communities rather than by isolated cells^19–21^. We postulated that while neighbourhood profiles may be conserved across patient TMEs, their prevalence and spatial organization vary across disease states. As such, we assigned cells to CNs based on the distribution of surrounding neighbouring cell types to represent the spatial TME architecture at the level of functional multicellular subcommunities [Fig.2a]. We identified 23 CNs with a prominent divide in malignant B-cell representation between CNs, featuring a B-cell predominant or non-B-cell predominant neighbour composition [Fig.2b]. Comparing the CNs between non-tFL and tFL-FL patients, the tFL-FL TME was composed of a significantly lower proportion of non-B_T-CM and non-B_Macro CNs than non-tFL [Fig.2d-e, Supplementary Fig.2a]. The non-tFL state displayed a contiguous and clustered distribution of non-B_T-CM and non-B_Macro cells, whereas these populations appeared more sparse and fragmented in tFL-FL (p = 0.036 from per-patient Wilcoxon rank-sum test [Fig.2d]. Cell-type enrichment analysis suggested a reduced enrichment of TFH, FDC, and DC populations in tFL-FL, with relative increases in macrophage, plasma, endothelial, and fibroblast compartments compared to non-tFL [Fig.2f].

**Fig 2.**
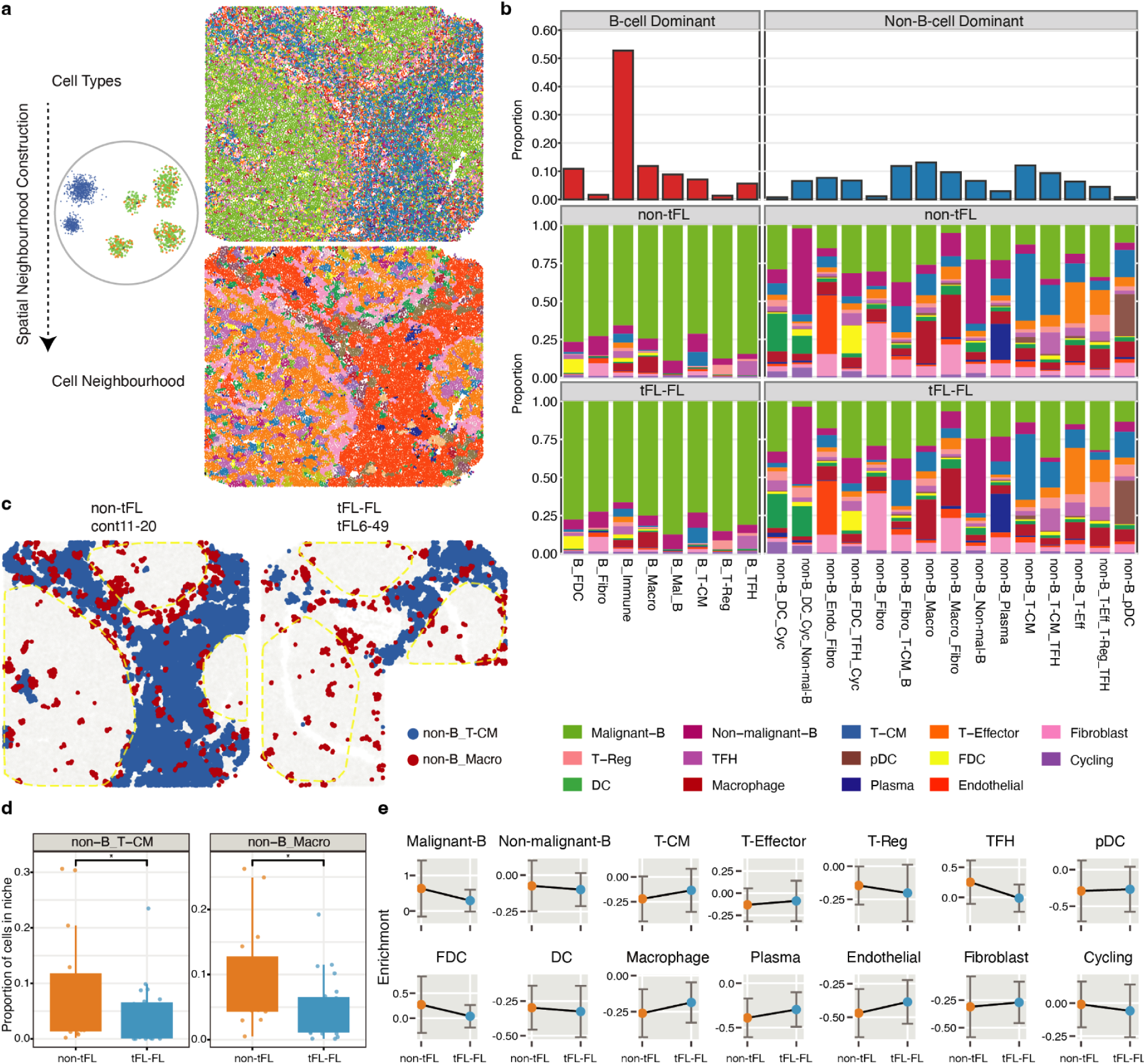
Recurrent cell neighbourhoods in the FL TME across disease states. **a,** Overview of recurrent cell neighbourhood construction. **b,** The cell neighbourhood composition in the B-cell-predominant and non-B-cell-predominant metacluster with stacked barplots of cell neighbourhood frequencies across all non-tFL and tFL-FL samples. **c,** Spatial mapping of cellular neighbourhoods in exemplar tissue cores, colored by selected cellular neighbourhoods. **d,** Boxplot showing the distribution of cell neighbourhoods of non-B_T-CM and non-B_Macro in non-tFL and tFL-FL (*: p<0.05; yellow dotted lines represent the follicular structures). **e,** Edge trend plot indicating enrichment of proximal inter-cell type interactions with respect to malignant B-cells in non-tFL and tFL-FL.

We next sought to investigate whether germinal centre (GC)-related expression in malignant B-cells was modulated by their specific neighbourhood context. We calculated a GC score for each malignant B-cell based on the expression of canonical GC-related gene signatures. Interestingly, the GC signature within the B_TFH CN was significantly stronger in non-tFL than tFL-FL [Supplementary Fig.2b]. This localized loss of GC identity suggests that a weakening of germinal center crosstalk occurs early in the pre-transformed TME, preceding overt architectural transformation. In contrast, GC scores within non-B-cell predominant CNs showed no substantial distinction across disease states [Supplementary Fig.2b]. Altogether, this indicates that FL transformation is characterized by shifts in cell abundance, but also a contraction of T-CM and macrophage-associated spatial neighbourhoods, which directly coincides with a localized loss of GC identity in malignant B cells.

### 3 Follicular architecture reveals divergent cellular and transcriptional programming in transformed FL

The malignant B-cell predominant follicular structure is a defining histological hallmark of FL and was observed in cores from non-tFL (n=6) and tFL-FL (n=9) via CosMx imaging. Cores taken from biopsies were arrayed on the TMA, with some cores containing large, medium, small, or no follicles, but all containing tumour cells. Utilizing pathology-guided computational mapping, we defined three spatial compartments: intra-follicular, peri-follicular, and extra-follicular zones [Fig.3a].

**Fig 3.**
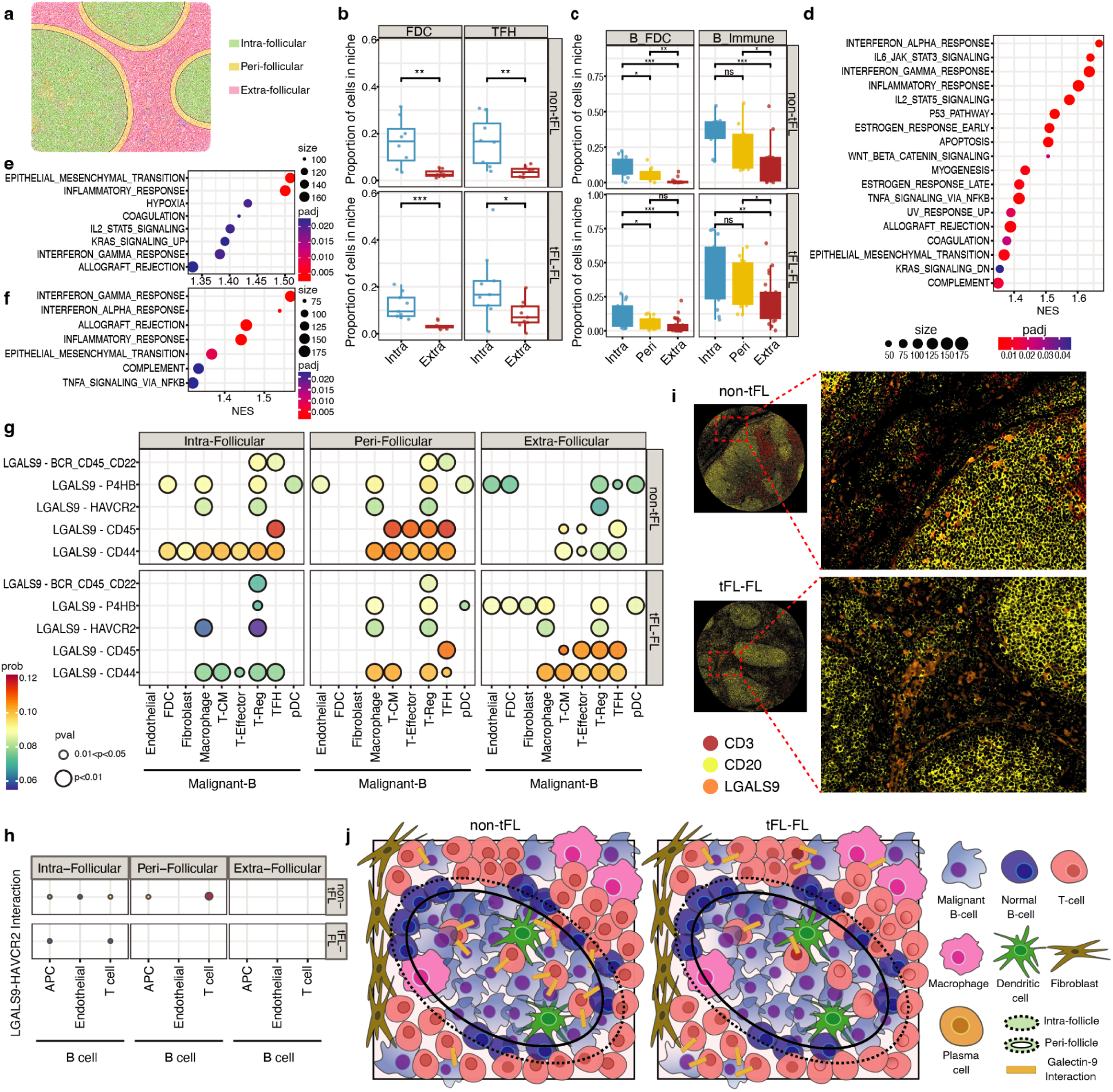
Spatiotemporal analysis of the FL TME across compartments and disease states. **a,** Spatial definition of intra-, peri-, and extra-follicular. **b,** Boxplot indicating the proportion of intra-follicular and extra-follicular FDC and TFH cells in non-tFL and tFL-FL disease states. **c,** Boxplot indicating the proportion of intra-follicular, peri-follicular, and extra-follicular B_FDC and B_Immune cell neighbourhoods in non-tFL and tFL-FL disease states. **d-f,** Gene set enrichment analysis showing normalized enrichment scores for significantly enriched pathways. In **d**, positive NES values indicate pathways enriched in peri-follicular zones compared to intra-follicular zones in non-tFL. In **e**, positive NES values indicate pathways enriched in peri-follicular zones compared to intra-follicular zones in tFL-FL. In **f**, positive NES values indicate pathways enriched in extra-follicular space compared to peri-follicular zones in tFL-FL. **g,** CellChat v2 analysis showing significantly enriched interactions within intra-follicular, peri-follicular, and extra-follicular zones of non-tFL and tFL-FL using CosMx data. **h,** CellChat v2 analysis showing significant LGALS9-HAVCR2 interactions between B-cells and microenvironment cells in intra-follicular and peri-follicular zones of non-tFL and tFL-FL using Cell DIVE data. **i,** Cell DIVE images of CD3 in dark orange, CD20 in yellow and LGALS9 in orange, in representative non-tFL and tFL-FL tissue cores. **j,** Model diagram comparing the spatial differences in LGALS9 signaling between non-tFL and tFL-FL.

We first assessed if compartmental composition and organization is disease state-specific. Consistent with the cellular organization of normal GCs, intra-follicular zones housed the majority of B cells, FDCs, TFHs, and T-Regs [Fig.3b and Supplementary Fig.3], while the rest of the TME cell populations were preferentially localized in extra-follicular space [Supplementary Fig.3]. This is also reflected in an enrichment of B_FDC and B_Immune CNs with a scarce amount of stromal CNs within the follicle [Fig.3c]. Although compartmental cell type composition was not significantly different between non-tFL and tFL-FL [Supplementary Fig.3], cell neighbourhoods exhibited compartment-specific localization and differed in spatial distribution between disease states. Specifically, B_TFH and B_T-Reg CNs were significantly enriched within the intra-follicular zone compared to the extra-follicular space of tFL-FL, whereas in non-tFL there was a much higher proportion of non-B_T-CM and non-B_FDC_TFH CNs [Supplementary Fig.4]. This supports the notion that the degree of intra-follicular FDC and TFH engagement with malignant B-cells may serve as a differentiating factor between non-tFL and tFL-FL.

As the peri-follicular zone is naturally the border of tumour-TME interplay, we were interested in how it differed to intra-follicular zones in neighbourhood composition. Interestingly, we observed primarily non-B-cell predominant CNs, including non-B_Endo_Fibro, non-B_Macro, non-B_Macro-Fibro, non-B_Plasma, non-B_T-CM, and non-B_T-Eff CNs having a significantly higher proportion in the peri-follicular zone in tFL-FL while the same effect was not observed in non-tFL [Supplementary Fig.4]. This suggests that unlike the indolent non-tFL, the tFL-FL state displays a greater localized accumulation of specialized non-B-cell predominant CNs at the immediate follicular boundary, highlighting the peri-follicular zone as a spatial domain of microenvironmental reprogramming.

We next examined signaling pathway heterogeneity across spatial compartments. In non-tFL, peri-follicular malignant B-cells exhibited a significant enrichment of immune activation and cytokine signaling pathways, including IFN-α and IFN-γ response, inflammatory response, and IL6-JAK-STAT3 signaling compared to their intra-follicular counterparts [Fig.3d]. Additionally, the peri-follicular malignant B-cells exhibited upregulated inflammatory, IFN-α response, and hypoxia activity compared to the extra-follicular space [Fig.3e]. In tFL-FL, malignant B-cells underwent a more profound spatial reprogramming between its compartments. Beyond immune cascades (interferons, TNF-α, complement), the peri-follicular zone also activated stress and apoptotic programs (p53, apoptosis, hypoxia), suggesting a heightened level of TME-mediated pressure near the boundary compared to within the follicle [Fig.3f].

We continued to investigate by comparing spatial compartments between disease states. The intra-follicular zone of tFL-FL were characterized by upregulated metabolic and proliferative activity among malignant-B cells, including MYC targets V1, oxidative phosphorylation, mitotic spindle, and G2M checkpoint pathways [Supplementary Fig.5a]. In contrast, intra-follicular malignant B cells in non-tFL were enriched for gene sets associated with myogenesis and angiogenesis, alongside a higher expression of *LGALS9* (Galectin-9) [Supplementary Fig.5b]. This transcriptional profile suggests active engagement with tissue remodeling to maintain structural homeostasis within follicles of a non-transforming TME^22^. The difference between the extra-follicular zones were minimal, with hypoxia being the sole enriched pathway in non-tFL compared to tFL-FL [Supplementary Fig.5c].

To comprehensively quantify cellular interactions across compartments with respect to disease states, we performed spatially-aware inference of intercellular communication that identified a striking shift in the *LGALS9*-related signaling axes [Fig.3g]. In non-tFL, *LGALS9*-related interactions were largely restricted to the follicle, while a nearly flipped pattern in tFL-FL was observed with signaling being exclusively significant in the non-follicular zones [Fig.3g]. This spatial shift was subsequently validated using Cell DIVE for measuring Galectin-9 and TIM3 intensity and distribution at the protein level [Fig.3h and 3i]. For each TIM3+ T cell, the proportion of neighbouring malignant B-cells expressing LGALS9 was significantly higher in non-tFL than in tFL-FL in both the intra-follicular and peri-follicular compartments (Mann-Whitney p = 0.044 and 0.031, respectively). Collectively, these compartmental analyses highlight that early FL transformation undergoes spatial reprogramming marked by heightened signaling at the peri-follicular boundary accompanied with a shift in LGALS9 signaling away from the follicular core [Fig.3j].

### 4 Macrophages exhibit differential polarization across follicular compartments and disease states

As neighborhood and compartmental analyses both highlight macrophage-associated spatial neighbourhoods as a feature of diverging TME dynamics between disease states, we were interested in further characterizing how macrophage populations varied in transcriptional programming with disease states and transformation.

Compared to tFL-FL, macrophages in tFL-DLBCL showed upregulation of TGF-β signaling and mitotic spindle pathways, suggesting enhanced pro-tumourigenic signaling and proliferative activity [Fig.4a]. Conversely, a broad spectrum of metabolic and cell cycle programs, including MYC targets V1, fatty acid metabolism, G2M checkpoint, coagulation, glycolysis, mTORC1 signaling, and E2F targets were significantly downregulated in tFL-DLBCL macrophages [Fig.4a]. This transcriptomic profile is directionally consistent with an alternatively activated, immunosuppressive tumour-associated macrophage (TAM) phenotype, leaning towards M2-like polarization^23,24^. In contrast, no significant transcriptomic differences were detected when comparing macrophages between non-tFL and tFL-FL, suggesting that their state remains relatively stable during the pre-transformation stage.

**Fig 4.**
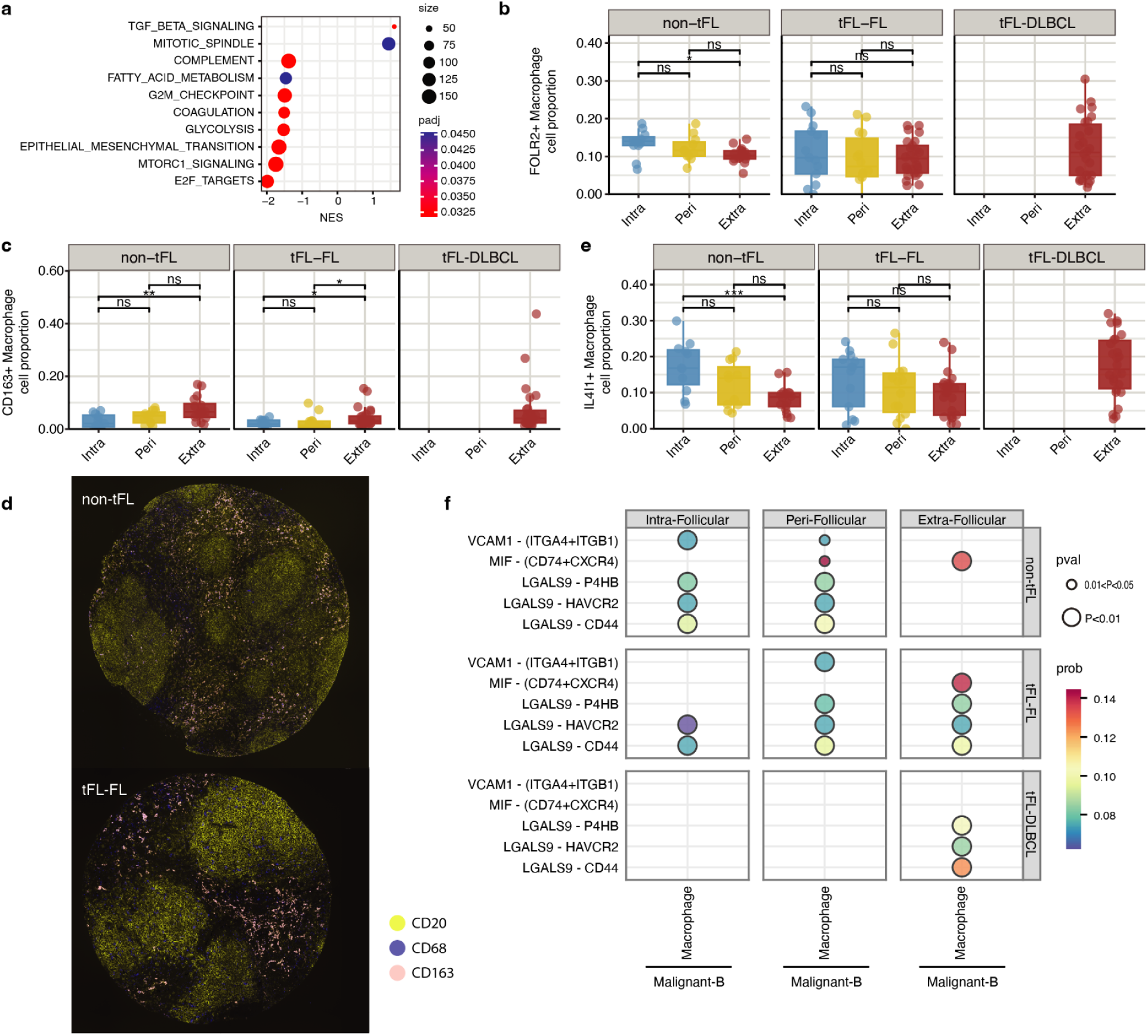
Analysis of macrophages in the FL TME. **a,** Gene set enrichment analysis showing normalized enrichment scores for significantly enriched pathways. Positive NES values indicate pathways enriched in tFL-DLBCL compared to tFL-FL. **b,** Boxplot indicating FOLR2+ macrophage proportion in intra-follicular, peri-follicular, and extra-follicular zones of non-tFL, tFL-FL, and tFL-DLBCL. **c,** Boxplot indicating CD163+ macrophage proportion in intra-follicular, peri-follicular, and extra-follicular zones of non-tFL, tFL-FL, and tFL-DLBCL. **d,** Cell DIVE images of CD20 in yellow, CD68 in blue, and CD163 in pink in representative non-tFL and tFL-FL tissue cores. **e,** Boxplot indicating IL4I1+ macrophage proportion in intra-follicular, peri-follicular, and extra-follicular zones of non-tFL, tFL-FL, and tFL-DLBCL. **f,** CellChat v2 analysis showing significantly enriched interactions within intra-follicular, peri-follicular, and extra-follicular zones of non-tFL, tFL-FL, and tFL-DLBCL using CosMx data.

In addition to gene set differences, we observed differences in the spatial distribution of specific macrophage subsets. In non-tFL, FOLR2⁺ tissue-resident macrophages (TRMs) were concentrated within or adjacent to follicles (p<0.02, Wilcoxon rank-sum test; intra-follicular vs. extra-follicular) [Fig.4b]. In both non-tFL and tFL-FL, CD163⁺ TAMs were relatively enriched in extra-follicular space (non-tFL p<0.01, tFL-FL p<0.05, Wilcoxon rank-sum test; intra-follicular vs. extra-follicular) [Fig.4c]. This spatial pattern was validated at the protein level for CD68 and CD163 via Cell DIVE [Fig.4d]. IL4I1⁺ macrophages also showed a bias for intra-follicular localization in non-tFL (p<0.01, Wilcoxon rank-sum test; intra-follicular vs. extra-follicular) [Fig.4e].

In tFL-DLBCL, the architectural organization of these niches collapsed as compartmental zones were absent in DLBCL morphology. CD163⁺ and IL4I1⁺ macrophage proportions were similar to those in the extra-follicular zone seen in non-transformed states, while FOLR2⁺ TRMs were similar to the intra-follicular and peri-follicular zones [Fig.4b-d]. These patterns mirror niche-resolved TAM architectures previously described^25^, suggesting that the loss of intra-follicular TRMs and the expansion of extra-follicular CD163⁺ macrophages in the zone represent a critical pro-tumourigenic remodeling of the TME.

Spatially-aware inference of cell-cell communication revealed that crosstalk between malignant B-cells and macrophages is both compartmentalized and disease state dependent. In non-tFL, interactions were pervasive across compartments. For instance, the VCAM1–(ITGA4+ITGB1) axis was detected in both intra-follicular and peri-follicular zones, while the MIF–(CD74+CXCR4) axis was mainly localized in extra-follicular space. In tFL-FL, however, these interactions became spatially displaced. VCAM1 signaling was absent within the follicle but increased in the peri-follicular zone and MIF signaling became exclusively extra-follicular [Fig.4f]. Interestingly, LGALS9-related pathways which were restricted to the intra-follicular and peri-follicular zones in non-tFL shifted to extra-follicular space in tFL-FL with an increased signaling probabilities. Overall, these results support a model where the crosstalk between macrophages and malignant B-cells is progressively displaced from the follicular zone to extra-follicular space in tFL-FL, accompanied by intensified LGALS9-mediated immunoregulatory signaling.

### 5 Transformation to DLBCL reshapes the spatial architecture of the tumour microenvironment

With matched pre-and post-transformation spatial measurements, we next sought to characterize the spatial microenvironmental reorganization accompanying disease transformation. We first observed amplified interactions between malignant B-cells and the TME from tFL-FL to tFL-DLBCL. Spatial graph modeling showed that the architectural shift to a diffuse state expands heterotypic interacting edges that are potentially associated with active TME remodeling. In contrast and as expected, FDCs and TFHs were sharply decreased in tFL-DLBCL [Fig.5a and 5b], reflecting the transition from preserved follicles in non-tFL and tFL-FL to the effacement of these structures in tFL-DLBCL. Aside from TFHs and FDCs, the proportion of T-CM cells exhibited a marked decline post-transformation, whereas T-effector, T-regulatory cells, and macrophages showed modest overall increases compared to the their pre-transformed counterparts [Fig.5b], consistent with observations made from matched scRNA-seq data^13^.

**Fig 5.**
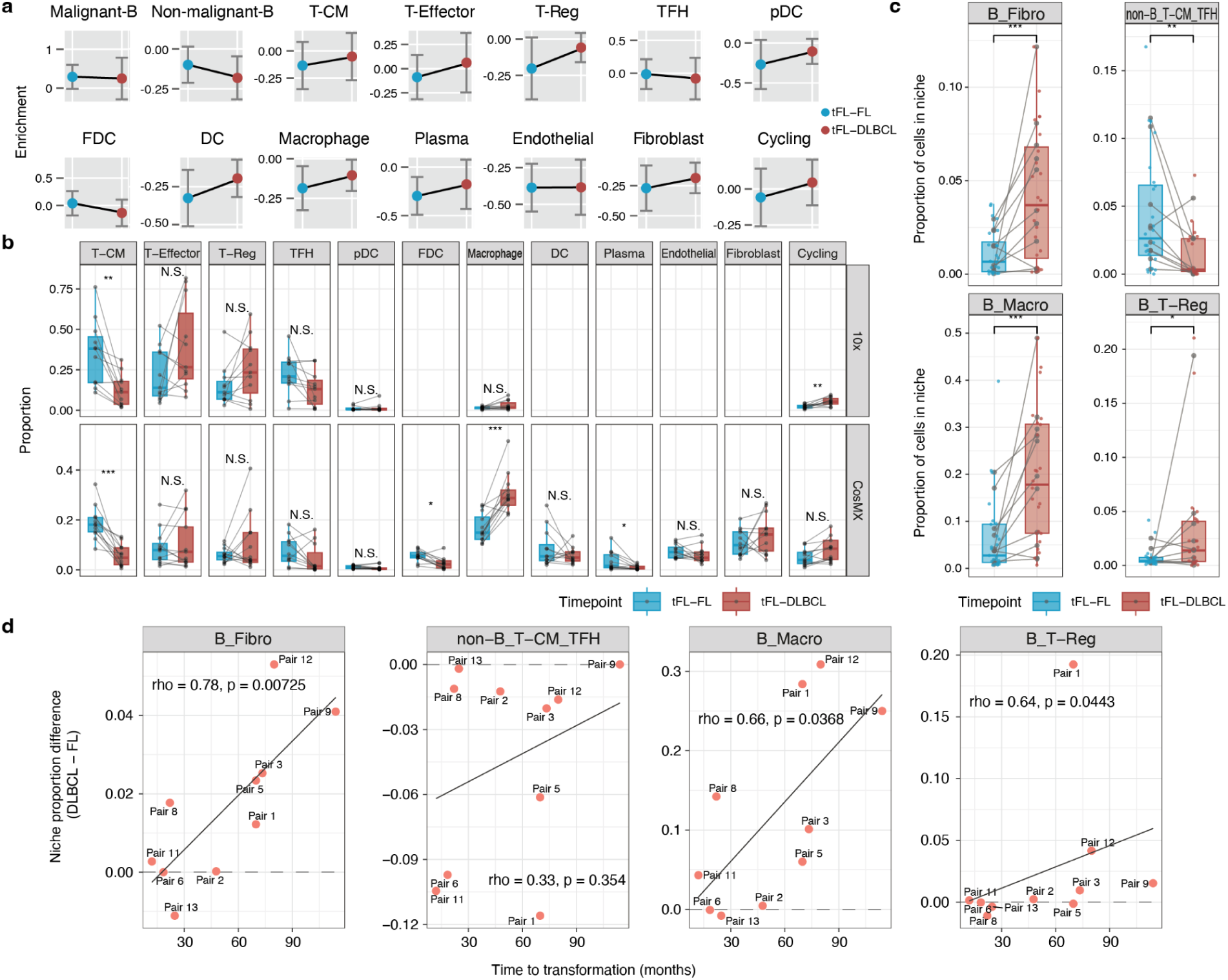
Spatial correlation with transformation to DLBCL. **a,** Spatial proximity between malignant B-cells and TME cell types across disease states. Cell-type enrichment around malignant B-cells was compared between tFL-FL and tFL-DLBCL samples. **b,** Paired comparison of cell-type proportions between tFL-FL and tFL-DLBCL in both the 10x and CosMX datasets. **c,** Changes in cellular neighbourhood prevalence after transformation. Selected cellular neighbourhoods were quantified in paired tFL-FL and tFL-DLBCL samples. Points represent individual samples and lines connect matched patient pairs (*: p<0.05; **: p<0.01; ***: p<0.001). **d,** Spearman association between time to transformation and paired changes in outside-follicular cellular neighbourhoods. For each patient pair, the proportional difference in each cellular neighbourhood was calculated as tFL-DLBCL minus tFL-FL and plotted against time to transformation in months.

We next assessed differences in CN prevalence with disease transformation. Compared to tFL-FL, tFL-DLBCL exhibited a significant increase in B-cell predominant CNs including B_Fibro, B_Macro, B_T-Reg CNs, alongside a decrease in non-B_T-CM_TFH CNs [Fig.5c]. The diffuse architecture of DLBCL permits malignant cells to escape follicular constraints and infiltrate stromal niches, where this shift favors a B-cell predominant neighbourhood landscape in the TME, for instance B_Fibro CNs are abundant in tFL-DLBCL while rare in tFL-FL [Fig.5c]. The reduction in non-B-cell predominant neighborhoods also likely reflects the expansion of DLBCL as dense sheets of malignant cells with fewer areas of stromal/inflammatory zones, which shift the TME from heterotypical interactions toward malignant B-cell-dominated niches.

We then examined whether the magnitude of CN remodeling was associated with the time to transformation. For each patient, the proportional difference for each CN was calculated as the difference between tFL-DLBCL and tFL-FL correlated with the interval to transformation. The result showed that the increase in B_Fibro CNs was positively correlated with time to transformation, indicating that cases with longer transformation intervals tended to show a larger gain of fibroblast-associated B-cell CNs [Fig.5d]. A similar positive association was observed for B_Macro and B_T-Reg CNs [Fig.5d]. In contrast, changes in non-B_T-CM_TFH showed positive but non-significant trends [Fig.5d]. These findings suggest that the expansion of selected B-cell-predominant stromal and macrophage-associated CNs are significantly associated with transformation intervals, whereas other niche changes are less strongly associated with transformation time.

In summary, spatial profiling of matched pre-and post-transformation pairs demonstrates that histological transformation exhibits a shift toward B-cell-predominant stromal and macrophage-associated CNs, in which the expansion of B_Fibro, B_Macro, and B_T-Reg neighborhoods positively correlates with the time to transformation.

### 6 Microenvironment remodeling in the DLBCL tumour microenvironment is associated with genome instability in malignant-B cells

Previous integrated single-cell whole genome sequencing (scWGS) of this cohort has suggested that genomic alterations partially underlie phenotypic transitions associated with transformation. Therefore, we here investigated whether specific spatial patterns were associated with elevated genome instability in malignant cells by dichotomizing patients by copy number alteration (CNA) thresholds. This resulted in a high-instability group of 5 tFL-DLBCL samples and 1 tFL-FL sample and a relatively stable group of 8 tFL-DLBCL samples and 11 tFL-FL samples [Supplementary Fig.5d]. To maintain statistical power, we focused our subsequent study on the tFL-DLBCL samples.

Comparison of gene sets between high-instability and relatively-stable groups revealed significant enrichment of proliferative and stress-related pathways, including E2F targets, MYC targets V1, G2M checkpoint, mitotic spindle, and oxidative phosphorylation pathways in genomic high-instability groups [Fig.6a]. Additionally, pathways involved in DNA repair and mTORC1 signaling were enriched [Fig.6a], consistent with the activation of genome maintenance and growth promoting programs in response to persistent DNA damage.

**Fig 6.**
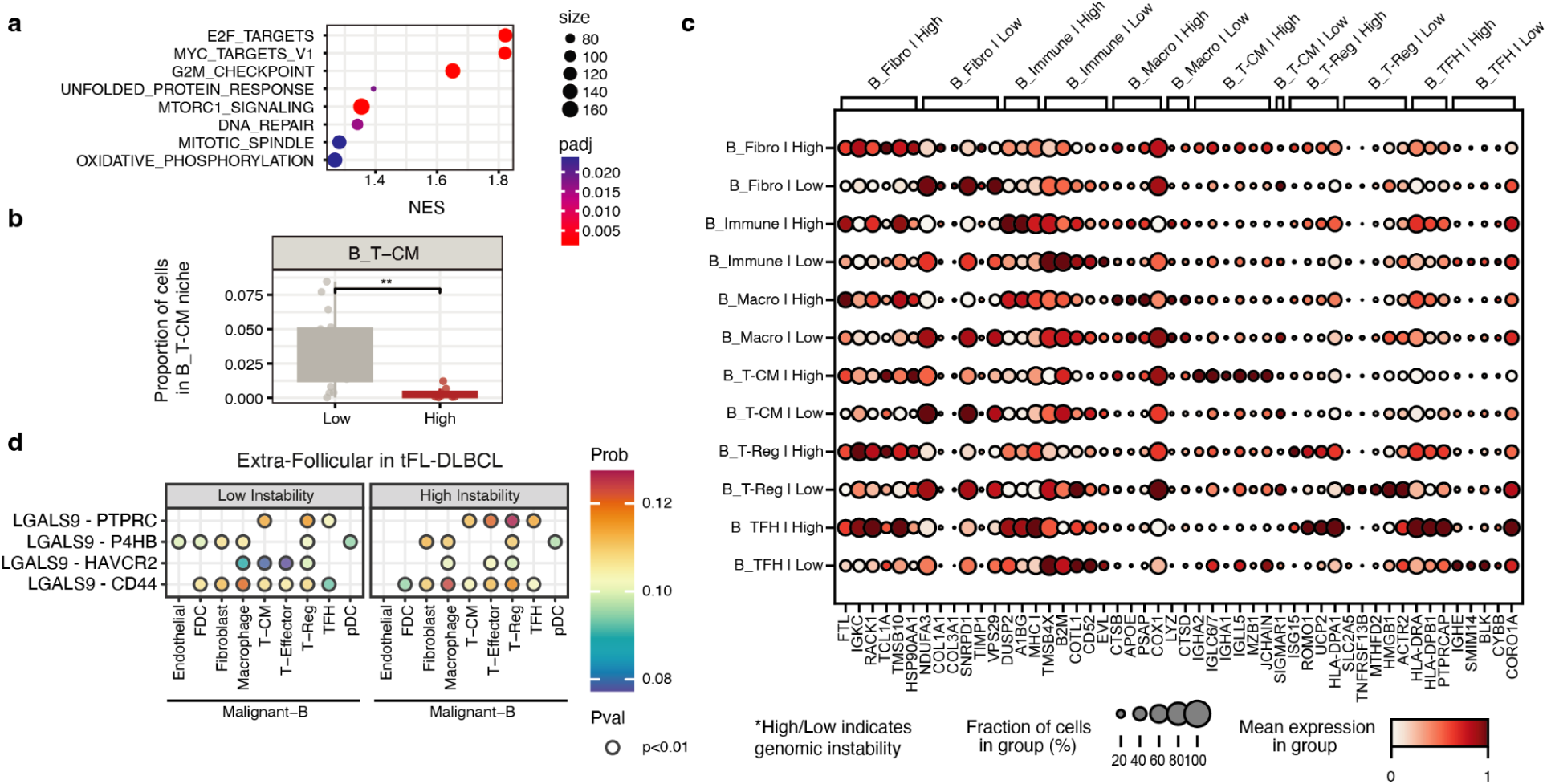
Analysis of TME phenotypic dynamics under differing genomic stabilities. **a,** Gene set enrichment analysis showing normalized enrichment scores for significantly enriched pathways. Positive NES values indicate pathways enriched in the genomic high-instability group compared to the relatively-stable group. **b,** Boxplot indicating proportion of genomic high-instability and relatively-stable cells in the B_T-CM cell neighbourhood. **c,** Dotplot representing the mean expression of transcripts in cell neighbourhoods dichotomized by high-instability and relatively-stable groupings. **d,** CellChat v2 analysis showing significantly enriched interactions within high-instability and relatively-stable groups in tFL-DLBCL.

While genomic instability in the malignant cells did not significantly alter the global proportions of TME cell types [Supplementary Fig.5e], spatial neighborhood analysis revealed a selective disruption in B–T-CM CNs in the high-instability group [Fig.6b and Supplementary Fig.5e]. This reduction points to a potential immune microenvironment reorganization that may impair supportive or regulatory immune systems. Highly variable gene analysis within CNs, however, illustrated a shift toward a more inflamed and antigen-presenting state in the high-instability group. Specifically, B_Fibro CNs displayed marked increases in extracellular matrix (ECM) remodeling genes (e.g. *COL1A1*, *COL3A1*, *TIMP1*, *MMP1*) [Fig.6c], suggesting an activated, fibrotic stromal program in unstable tumours. Pan-immune (B_Immune) and macrophage (B_Macro) predominant CNs were characterized by increased expression of lysosomal and phagocytic markers (e.g. *LYZ*, *CTSB*, *CTSD*, *APOE*, *PSAP*, *CYBB*, *CORO1A*), and MHC class II molecules (*HLA-DPA1*, *HLA-DRA*, *HLA-DPB1*) [Fig.6c], indicating an association between genomic instability and enhanced antigen presentation and immune activation. For T-cell–rich CNs (B_T-CM, B_T-Reg, B_TFH), acquired inflammatory and interferon-response signatures (e.g. *ISG15*, *HMGB1*, *ROMO1*, *UCP2*) [Fig.6c], further suggesting that even T cell–dominated structures adopt an activated state while residing within TME associated with high-instability tumour contexts.

We further probed at whether genomic instability drives differential spatial interaction patterns. Samples in the relatively-stable group exhibited a higher number of interactions from malignant B-cells to endothelial cells, fibroblasts, macrophages and multiple T-cell subsets [Fig.6d]. In contrast, high-instability tumours were characterized by a selective accentuation of LGALS9 signaling (*LGALS9–HAVCR2*, *LGALS9–CD44* and *LGALS9–P4HB*) from malignant B-cells to T-Reg, T-Effector, and TFH cells [Fig.6d]. Chemokine and adhesion pathways (*CCL17–CCR4/ACKR1*, *CXCL13–CXCR5/CXCR3*) were also significantly stronger, suggesting increased recruitment and retention of immune and stromal constituents [Fig.6d]. Together, this suggests that high genome instability is associated with a rewiring of malignant B-cell communication, featuring immunoregulatory crosstalk, chemokine-mediated trafficking, and intensified adhesive signaling within the tFL-DLBCL TME.

## Discussion

We mapped the spatial architecture of follicular lymphoma and defined the cellular and molecular dynamics underlying disease transformation through integrated single-cell spatial omics. By profiling 10 non-transformers and 12 transformers with paired biopsies, we phenotyped individual cells, quantified the interactional structures of neighborhoods and compartments, and correlated these features with genomic stability. This multi-layered analysis shows that FL transformation is accompanied by the progressive disappearance of organized, FDC-and TFH-supported follicular niches and a coordinated shift toward a diffuse, stromal-rich and immunosuppressive microenvironment, including polarization of macrophages toward CD163+/IL4I1+ immunosuppressive states, repositioning of Galectin-9 and TIM3 interactions out of the follicle, and a reorganization of malignant B-cell neighbourhoods linked to genomic instability.

Our findings demonstrate that compartmental composition and cell neighbourhood organization are disease state-specific. While scRNA-seq analyses have previously suggested transformation is dominated by a change in the composition of T cell subsets^13^, spatial transcriptomics reveals that these shifts are driven by the emergence of specific cellular neighbourhoods. We observed that in the transition from tFL-FL to tFL-DLBCL, the TME shifts from a heterotypical architecture toward one dominated by malignant stromal-heavy B-cell-predominant neighborhoods. This remodeling is characterized by the effacement of follicles and the induction of reactive stromal programs, providing a structural basis for the aggressive growth of transformed cells.

Our compartment-resolved transcriptional analysis further suggests that the peri-follicular region may represent a critical transition zone during disease progression. As malignant cells exit the follicular compartment, they retain inflammatory and stromal-interactive signatures while showing a reduced apoptotic profile, consistent with stabilization in an invasive, TME-reactive state. Compared with non-tFL, the tFL-FL state exhibits a broader and more pronounced spectrum of stress-adaptive and stromal-interactive pathways, indicating that transcriptional remodeling is already evident before overt histologic transformation. This divergence between disease states suggests that FL transformation is accompanied by compartmental remodeling of spatial signaling dynamics, with important implications for malignant B-cell interaction with the surrounding TME.

Central to this spatial reorganization is the spatial displacement of macrophage subsets. While FOLR2⁺ TRMs remain sequestered within follicles in non-progressors, the transformation process is marked by their depletion and the concurrent expansion of CD163⁺ and IL4I1⁺ immunosuppressive TAMs in the extra-follicular space. This spatial polarization toward an M2-like phenotype suggests that the TME in the pre-transformed state is potentially primed to support immune evasion.

Our integration of whole-genome sequencing highlights genomic instability as a key feature associated with this TME rewiring. In highly unstable tFL-DLBCL samples, we observed a selective disruption of supportive B–T-CM neighborhoods and a sharp upregulation of fibrotic and metabolic programs. These tumours exhibit a stressed but highly proliferative phenotype, characterized by intensified chemokine-mediated recruitment and adhesive signaling. This suggests that genomic instability does not just fuel intrinsic malignancy but actively shapes a more inflamed and stromal-rich TME that may be more resistant to conventional therapies.

Perhaps the most captivating finding from our spatially-aware crosstalk analysis is that *LGALS9* signaling exhibits differential compartmental localization between non-tFL and tFL-FL. Galectin-9 is a β-galactoside lectin protein that can crosslink glycoproteins to regulate cellular processes via multiple ligand-receptor interactions^26^. It is known to interact with PD-1 and TIM-3 to regulate T cell senescence and inactivation. Notably, anti-galectin therapy can block this interaction, thereby allowing CD8⁺ T cells to potentiate anti-tumour immunity^27^. Galectin-9 also interacts with leukocytes to regulate B-cell activation within germinal centers^28^. Interestingly, we observed that non-tFL specimens contained Galectin-9 expressing malignant B-cells interacting with TIM-3 positive T cells in intra-follicular and peri-follicular zones, whereas this interaction shifted to the extra-follicular space in pre-transformation FL tumours. Given that non-tFL follicles display stronger germinal center gene signatures, it can be hypothesized that Galectin-9 is able to engage in interactions facilitating normal regulatory functions in the follicular core of non-transformers, while these interactions transform to be pro-tumourigenic and immunosuppressive in the pre-transformed TME as germinal center signatures decline. Rather than global overexpression, the spatial localization of Galectin-9 signaling appears to differentially modulate disease transformation. A recent study on patient-derived follicular lymphoma spheroids has shown evidence that anti-Galectin-9 treatment improves rituximab efficacy, highlighting Galectin-9 as a promising emerging therapeutic treatment strategy^29^.

We acknowledge several limitations to this study. The tissues of interest were embedded in TMAs, which limited sampling to tissue cores. While we analyzed two cores per tumour, this provides a relatively limited perspective of the total tumour volume. Future studies with a larger cohort with increased statistical power may further validate these findings. However, the robustness of our results are supported by the high degree of concordance across multiple orthogonal modalities, including spatial proteomics and single-cell whole transcriptome sequencing. Collectively, our work provides a high-resolution spatial map of FL transformation, identifying the peri-follicular zone and Galectin-9 signaling as critical focal points for future therapeutic intervention.

## Methods

### Single-cell spatial cohort

This study has been reviewed and approved by the University of British Columbia–BC Cancer Agency Research Ethics Board (H22-03661), in accordance with the Declaration of Helsinki. Written informed consent from the patients has been previously obtained or the need for consent was waived in this retrospective study. The single-cell spatial cohort overlaps with the single-cell discovery cohort described previously^13^, with additional non-transformed and transformed patient samples. Duplicate 1 mm formalin-fixed paraffin-embedded (FFPE) cores were obtained from representative regions of follicular lymphoma of 10 non-tFL and 12 tFL pairs with pre-transformation (tFL-FL) and post-transformation (tFL-DLBCL) timepoint biopsies, along with one additional transformed patient (tFL10) with a post-transformation biopsy only, to construct a tumour microarray (TMA). Two tonsil and liver tissue control cores respectively were also included.

### Tissue processing and data acquisition for CosMx

The SSC CosMx dataset was generated by using the CosMx Spatial Molecular Imager (SMI) instrument from Nanostring Technologies following the manufacturer’s protocols in the CosMx SMI Manual Slide Preparation for RNA Assays (MAN-10184-06). Briefly, a 5 µm tissue section from the FFPE TMA was placed onto a Superfrost Plus microscope slide. Deparaffinization, heat-induced epitope retrieval, and enzymatic permeabilization were performed. Fiducials were applied to the slide, which were then post-fixated and blocked using NHS-acetate. Hybridization was done using probes from a predefined customized panel. After washing out unbound probes, slides were stained with nuclear and segmentation markers (DAPI, CD45, CD68, CD3, CD298). To assemble the flow cell, a coverslip was placed on the slide and loaded on a fluidic manifold onto the instrument.

A maximum of four FOVs were chosen and reviewed for each core by a pathologist. All FOVs were then subjected to the instrument for spatially resolved RNA profiling using a predefined customized panel with in-situ hybridization (ISH) probes and fluorescent readout probes. The RNA panel consists of 6375 ISH probes, combining the CosMx Human 6K Discovery panel with an addition of 200 custom selected genes relating specifically to B-cell lymphomas. The details of the panel are listed in Supplementary Table 2. Probe fluorescent signals in the raw images were detected at subcellular resolution and aggregated for transformation into decoded RNA transcripts at their registered three-dimensional global spatial location. After examining tissue section quality, a total of 69 cores from 23 patients and 1 tonsil sample were profiled (1 tonsil: 2 cores; 10 non-tFL: 16 cores; 1 tFL-DLBCL from tFL10: 2 cores; 12 tFL pairs: 49 cores). The tFL-FL timepoint was removed for tFL10 due to poor tissue quality.

### Image processing, single-cell segmentation and filtering for CosMx

Pre-processing and downstream analysis described onward was performed using the spatial framework, Sakura^40^. Multiplexed immunofluorescent stainings of DAPI, CD45, CD68, CD3, and CD298 proteins were co-measured on the FOVs along with RNA quantification. Specifically, DAPI, CD45 and CD68 were used as inputs for cell segmentation. Single-cell masks were generated through a set of raw image pre-processing steps followed by a deep learning-based segmentation method, Mesmer^30^, implemented in Sakura. Various segmentation algorithms were considered, while the best performing was chosen after systematically comparing with other methods^31^, such as the manufacturer variant of Cellpose^32^. Transcripts were mapped onto single-cell masks to generate a cell by transcript count matrix. This gene expression matrix was then filtered to retain high-quality cells for downstream analysis based on the following criteria: (1) cell area > 0; (2) total number of transcripts > 20; (3) percentage of negative probe < 5%. FOVs were lastly stitched together to recover core samples based on the global position of each core.

### Normalization and cell phenotyping for CosMx

The resulting gene expression matrix was normalized by library size, followed by a log transformation. De novo unsupervised clustering was done using the Leiden algorithm^33^ on the dimensionally reduced gene expression matrix. Principal component analysis was performed on the whole gene expression matrix and the top 30 PCs were used as input. B-cell clusters and non-B/TME clusters were classified based on the gene expression of B-cell lineage markers within each cluster. To improve delineation within each subpopulation, the clusters were subjected to re-clustering.

Based on a disaggregated single-cell phenotyping approach, non-malignant B-cell clusters and malignant B-cell clusters were classified by using the patient entropy scores and kappa/lambda ratios. A patient entropy score was defined to assess patient heterogeneity and timepoint heterogeneity in each cluster. Clusters with high patient entropy were selected as potential non-malignant B-cells as non-malignant B-cells share similar expression profiles across samples. For malignant B-cells, low patient entropy was expected as malignant B-cells tend to be patient specific. Non-malignant B-cells also tend to have a 60:40 ratio of kappa and lambda light chain expression while this ratio is often altered to either almost complete kappa or almost complete lambda expression in malignant B-cell clusters. Final B-cell annotations were examined by expertise from pathology.

For non-B/TME clusters, the z-score median expressions of clusters were assessed to merge clusters with similar phenotypes. Subsequently, cell types were annotated based on the characteristic gene expression of known lineage markers.

### Cell neighbourhood annotation for CosMx

Recurrent cell neighbourhoods were computed for all cells using Sakura^40^. For each cell, its neighbourhood is defined as all neighbouring cells within 18 µm between cell centroids. The defined distance was evaluated by grid searching across various distances and evaluating results through Spearman correlation and neighbourhood phenotype distribution. The phenotype of a cell neighbourhood is then calculated based on the normalized cell type count distribution. All cell neighbourhoods were clustered using a two-step approach. Otsu thresholding was first applied to the neighbourhood to dichotomize all neighborhoods into B-cell dominant or non-B-cell dominant neighbourhoods. The two types of neighbourhoods were then clustered further to annotate granular clusters by using the Leiden algorithm^33^. For B-cell heavy neighbourhoods, large clusters were annotated first to train a random forest classifier to annotate the rest of the clusters by merging clusters to the most similar cluster.

### Follicular structure annotation

Follicular structures in all cores were annotated using Sakura^40^ and examined by a pathologist. Briefly, hematoxylin and eosin stains as well as multiplexed immunofluorescence stains were both used to determine regions of FOVs with follicles. Based on review, a convex hull enhanced density-based clustering method was then used to classify each cell as a cell in a follicle or a microenvironment cell. Cells in follicles were numbered according to their belonging follicle. Cores with tissue tear required manual adjustment to merge cells to the same follicle. Cells were also annotated in another layer as within the follicular boundary region if cells were positioned 18 µm within the follicle or 18 µm outside of the follicle.

### Germinal center signature score for CosMx

Gene signatures for germinal centers (GC) cells were adapted from previously published work^34^. The gene signature was restricted to genes that were captured in our panel, leaving a total of 32 genes. A GC score was then calculated from the gene signature using the score_genes function from Scanpy^35^, which subtracts the summed expression of all genes for a given signature by the average of those genes. The GC score was validated using a germinal center from a reactive lymph node.

### Differential gene expression and enrichment analysis for CosMx

Differential gene expression was performed using Monocle 3^36^ with covariate terms including timepoint, patient and spatial regions (intra-follicular, peri-follicular, or extra-follicular), genome instability. The resulting values were then used as input for gene set enrichment analysis with the pre-ranked mode in fgsea^37^, using default parameters with 1000 permutations, and hallmark pathway gene set (The Molecular Signatures Database Hallmark Gene Set Collection). For pathway enrichment, genes with a significant p-value were ranked by log fold change in differential expression between the non-tFL and tFL-FL. All reported Q-values refer to Benjamini-Hochberg corrected p-values for two-sided tests.

### Spatial enrichment and cellular interaction analysis for CosMx

To analyze the enrichment of spatially interacting neighbours, we computed an adjacency enrichment score defined in Sakura^40^. Briefly, the adjacency enrichment score is the ratio between the total pairs of adjacent cells of interest and the expectation of the number of pairs.

The expectation is computed as the two times the multiplication of the proportions of the two types of cells of interest by the total number of pairs across all cells, subtracted by 1.

To examine ligand-receptor interactions and secreted signaling, CellChat v2^38^ was employed to infer spatially proximal crosstalk between malignant B and TME cells. CellChat was run for cells in follicular, follicular boundary, and non-follicular spatial regions respectively for each timepoint (namely non-tFL, tFL-FL, and tFL-DLBCL). CellChat v2 analyzes intercellular communication pathways with spatial context with support from a database of interactions among ligands, receptors and their cofactors that represent known heterodimeric molecular complexes. The database was adapted to include the MHCII-LAG3 ligand-receptor pairs and the signaling pathway categories used were secreted signaling and cell-cell contact. Significant signaling pathways were selected via a permutation test.

### Genome instability annotations using single-cell whole genome sequencing

All transformed tFL-FL and tFL-DLBCL pairs previously sequenced using single-cell whole genome sequencing (DLP+)^39^ were annotated with high or low genome instability. High genome instability was annotated for samples with a proportion of non-diploid genomic bins larger than 25 percent, while the inverse being low genome instability. Genomic bins were defined as the non-overlapping 500,000 genomic regions over the genome and the reads from DLP+ data were tabulated for the bins. Copy number was computed using HMMcopy, fitting on 6 possible ploidy settings and the best fit was computed for each genomic bin of cells.

### Tissue processing and data acquisition for Cell DIVE

The SSC Cell DIVE dataset was generated by using the Cell DIVE multiplexed imaging solution instrument from Leica Microsystems. Briefly, a 4 µm tissue section from the FFPE TMA was placed onto Apex Bond adhesive slides (Leica, 3800040). The mounted tissue was dried for 24 hours in a fume hood at room temperature. Slides were then baked and dewaxed on the BOND Rx autonomic stainer (Leica, 21.2821) using manufacturer recommended settings, and a dispense setting of 150 µl for all steps. Antigen retrieval was carried out sequentially, first in BOND ER1 buffer (Leica, AR9961) for 20 min at 100 °C, and then in BOND ER2 solution (Leica, AR9640) for an additional 20 min at 100 °C. Slides were then permeabilized with 0.3% Triton PBS for 30 minutes followed by blocking with Antibody blocking solution (Akoya, ARD1001EA) for 1 hour. Lastly, slides were stained with 0.2 ng/mL DAPI stock solution (Thermofischer, D3571) prepared in 1xPBS for 10 minutes. Slides were then taken out of the BOND Rx stainer and washed with deionized water in a coplin jar.

The slide was then subjected to repeating cycles of antibody labeling, image acquisition, immunofluor cleavage, and slide washing. Whole slides were scanned at 20x on the Leica Cell DIVE instrument in 1X PBS with 50% glycerol (Sigma, G5516) following manufacturer recommended settings. After each autofluorescence cycle, tissues were stained with directly conjugated antibodies diluted in a cocktail in antibody blocking solution (Akoya, ARD1001EA) for 1 hr at room temperature with gentle shaking. Tissues were then washed 3×2 minutes with PBS Tween 0.01%, followed by 2-minute refresh in DAPI, then immediately imaged at 20x. Dye inactivation was done by incubating whole slides in 0.5 M sodium bicarbonate (Sigma, S5761) solution prepared in DI water with 3% hydrogen peroxide (Sigma, H1009) for 15 minutes at room temperature.

A predefined panel of 21 fluorescent-dye conjugated antibodies were prepared and designed to target major cell types in FL Supplementary Table 3. Image tile merging, registration, autofluorescence and illumination correction were processed by the instrument software. Raw tiff images were outputted, and the overall staining quality was examined by a pathologist.

### Computational processing, TME phenotyping, and spatial analysis for Cell DIVE

Computational pre-processing of Cell DIVE was done using the Aivia image analysis software from Leica Microsystems with downstream analysis done using Sakura^40^. Single cells were segmented using the multiplexed cell detection method in Aivia. The last round of DAPI nuclear staining and 9 membrane stains (*CD20, CD68, CD21, CD3, CD11c, CD8, CD4, CD31, CD38*) were used as input for segmentation. The mean signal intensity for each protein was measured for each cell, which was organized as a cell by protein signal matrix. To remove background noise from imaging and tiling, the signal intensity was clipped if under 10. The cell by protein matrix was log transformed to normalize signal distributions.

Cell types were first annotated by signal thresholding a subset of cells with the highest intensity of proteins indicative of cell lineage. In addition to log normalization, signals were z-score scaled for phenotyping. A random forest classifier was trained based on the normalized intensity profiles of annotated cells to annotate the rest of the cells.

Annotations of cell neighbourhoods and compartments (intra-follicular, peri-follicular, and extra-follicular) were done in similar fashion as described above for CosMx with modifications to the distance metric from 15 µm.

### Statistical analysis

Statistical analysis was done using packages from both Python and R. All statistical details including the statistical test, the number of samples, and (adjusted) p-values can be found in the results, figures, and figure legends. Statistical significance was defined as p-value less than 0.05. The Benjamini-Hochberg procedure was used for false discovery rate corrections on p-values. Wilcoxon rank-sum tests were used for pairwise comparisons unless otherwise stated.

## Data Availability

CosMx and CycIF to be submitted to Zenodo and will be available upon publication.

## Contributions

S.W., E.L., C.S., A.R.: project conception and study design. S.W., E.L.: manuscript writing and editing. S.W., E.L.: computational biology and data analysis. S.W., E.L., A.S.F., T.H.: data visualization. A.T., Y.Y., A.F., S.R., M.F., T.A., D.W.S., C.S.: CosMx data generation and pre-processing. E.L., S.D.M., H.C., D.G.M.: Cell DIVE data generation and pre-processing. A.L., S.D.M., S.A., T.A.: pathology review.

## Acknowledgements

We would like to give our thanks to the patients and clinical staff involved in this study. E.L. was awarded a Canada Graduate Research Scholarship – Doctoral (CGRS D) from the Canadian Institutes of Health Research. A.R. receives operating funds from the Natural Sciences and Engineering Research Council of Canada (grant RGPIN-2022–04378), Terry Fox Research Institute (grant 1108).

## Ethics Declaration

C.S. performed consulting for Bayer Inc. D.W.S. reports receiving honoraria from and serving on advisory boards for Arima Genomics, AstraZeneca, Chugai, Eli Lilly, Kite/Gilead, and Roche.

## Extended Data

**Extended Figure 1.**
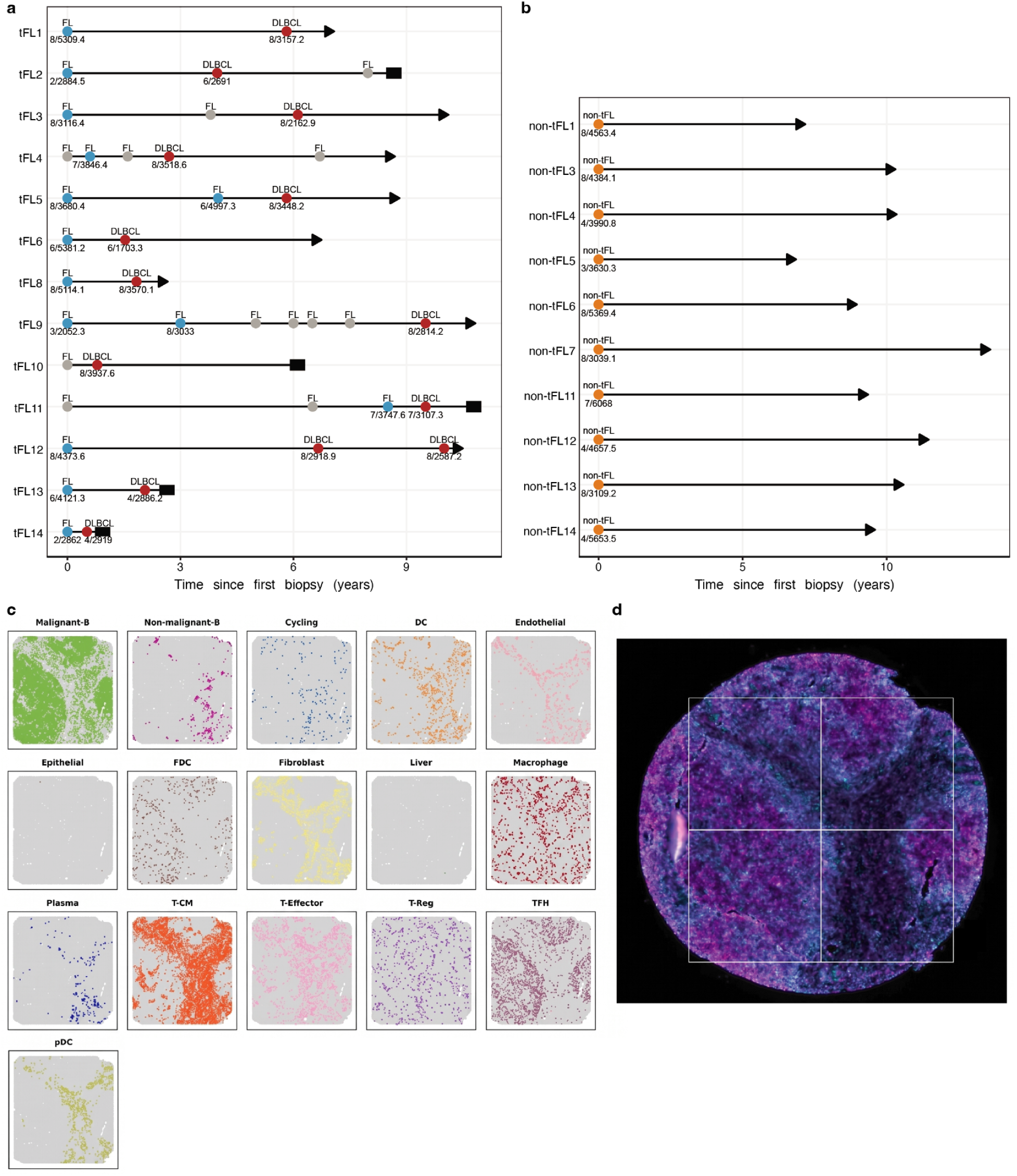
Overview of the cohorts and cell type annotation in spatial context. **a,** Clinical details of the paired cohort and each patient with two samples: tFL-FL (blue) and tFL-DLBCL (red). Of note, the tFL-FL sample is missing for tFL10. **b,** Clinical details of the non-tFL cohort (yellow). **c-d,** Example of cell type annotation in spatial context of non-tFL11_20 sample and the corresponding CD45 staining.

**Extended Figure 2.**
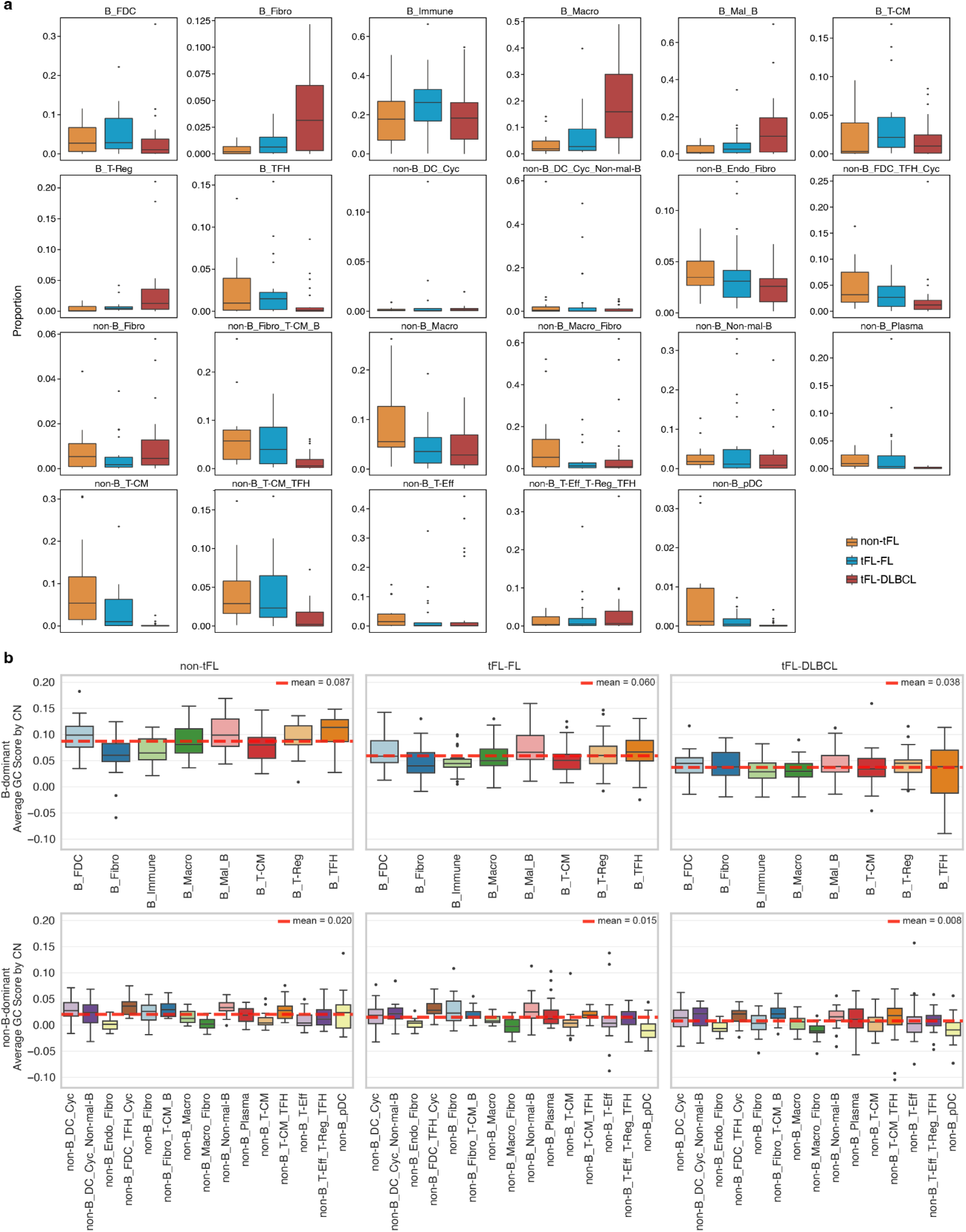
Cell-neighborhood prevalence comparisons across disease timepoints and GC-score patterns across B-dominant and non-B-dominant CNs and disease states. **a,** Boxplots showing the per-sample proportion of selected cell neighbourhoods (CNs) across disease states (non-tFL, tFL-FL, tFL-DLBCL). Each point represents one sample; boxes summarize the distribution across samples. **b,** Boxplots of GC scores for malignant B-cells stratified by different CNs and disease state. Distributions are shown per CN with disease-state– specific colouring.

**Extended Figure 3.**
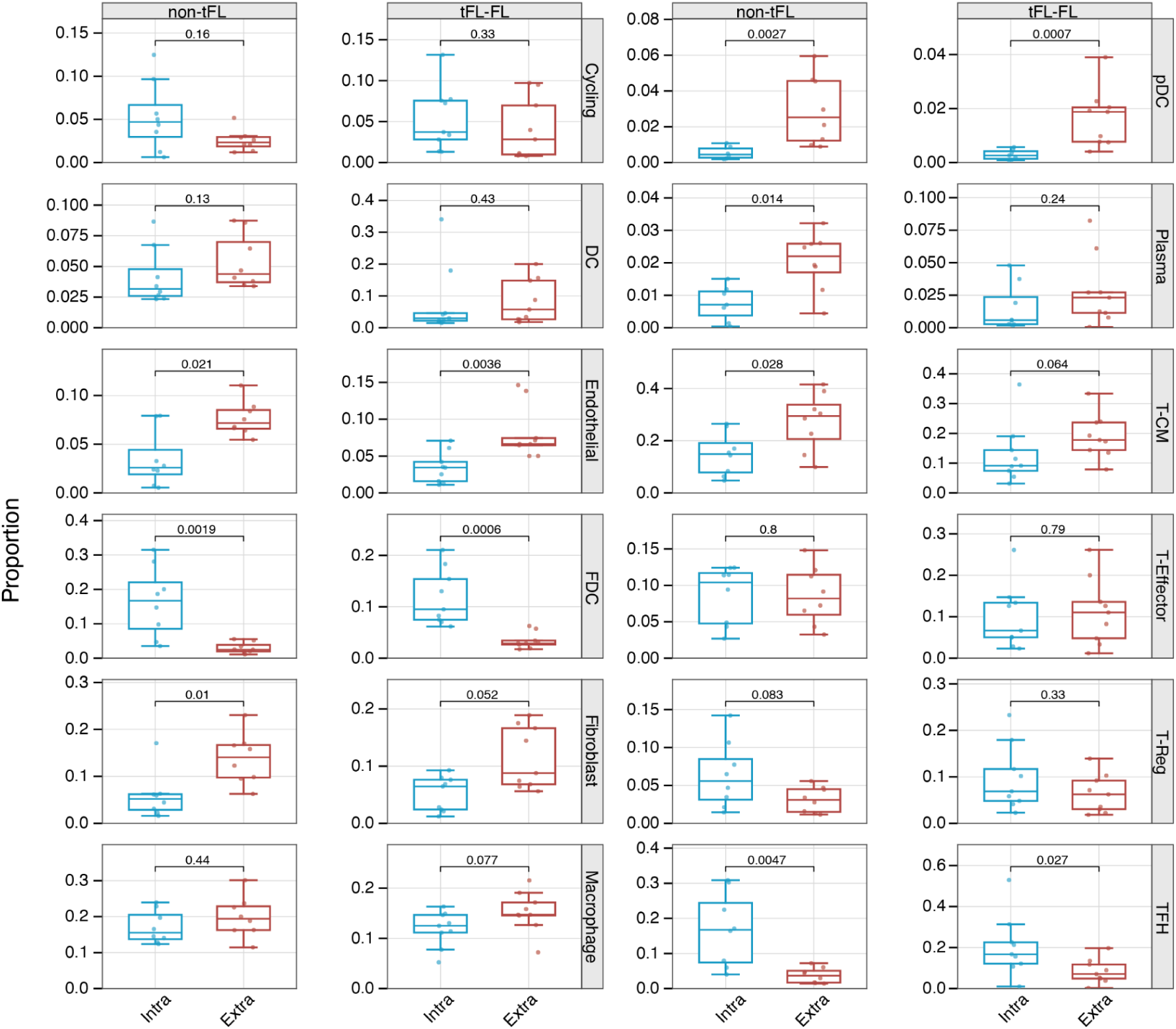
Spatial cell-type composition across follicular compartments. Barplots summarizing cell-type proportions across spatial compartments (intra-follicular vs extra-follicular regions), stratified by disease group (non-tFL vs tFL-FL).

**Extended Figure 4.**
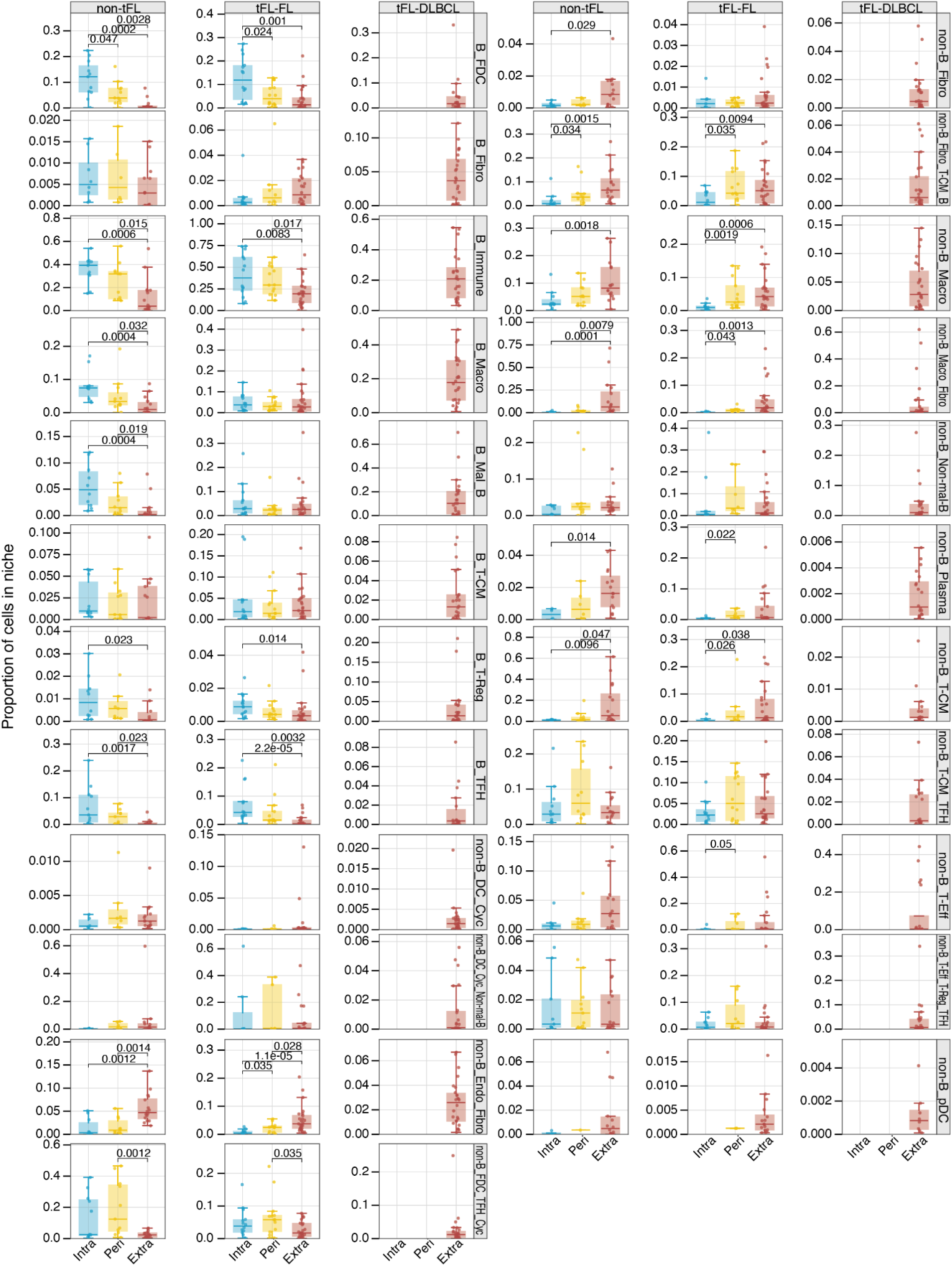
CN distribution by spatial compartment within timepoints.

**Extended Figure 5.**
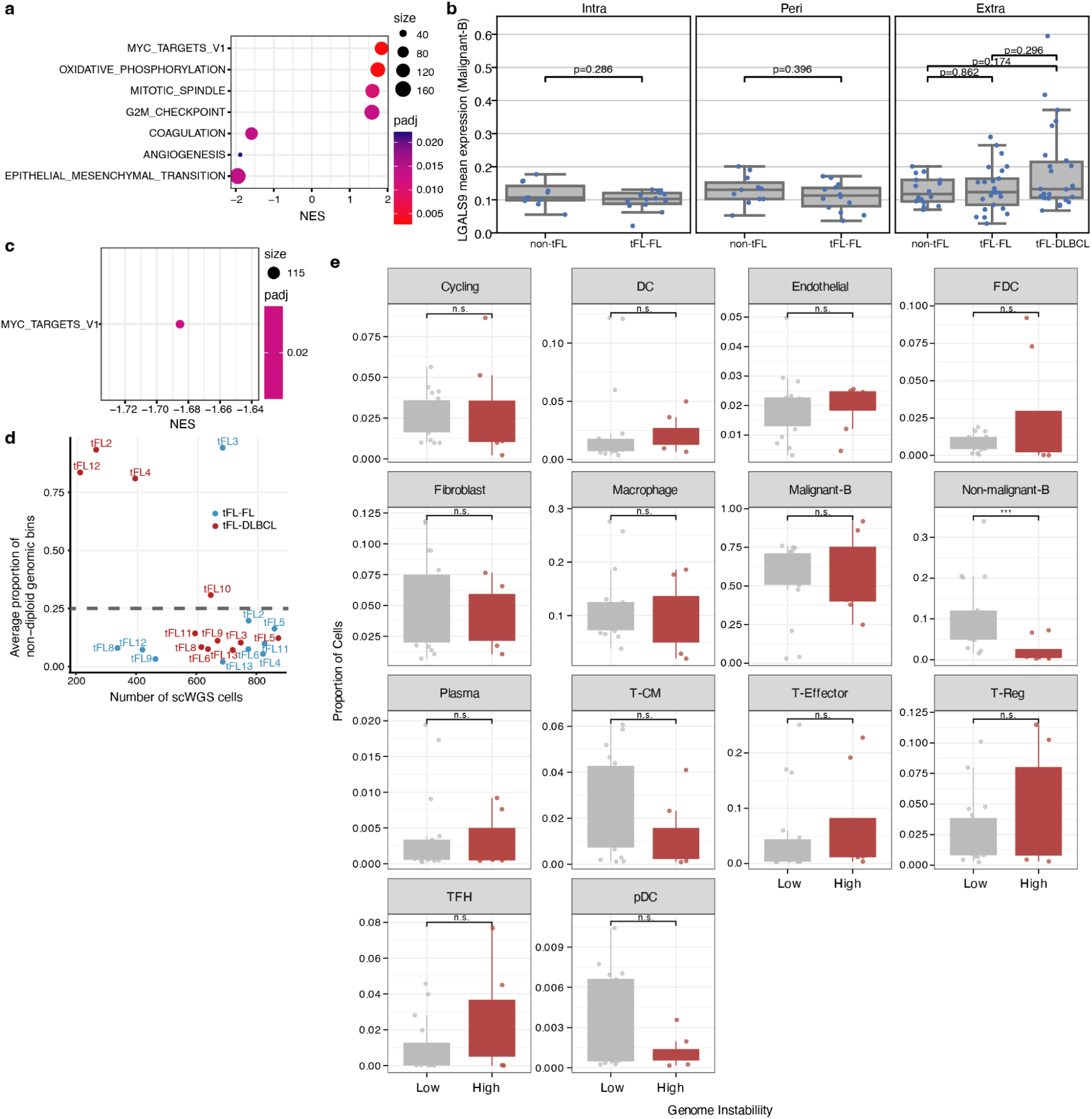
Compartment-resolved transcriptional programs and genome-instability characterization. **a,** GSEA result of intra-follicular regions between non-tFL and tFL-FL. The x-axis shows the normalized enrichment score (NES). **b,** Spatial pattern of LGALS9 expression in malignant B-cells. Boxplots of LGALS9 (Galectin-9) expression in malignant B-cells across disease states and spatial compartments (intra-, peri-, extra-follicular regions). **c,** Hypoxia enrichment in the extra-follicular region in tFL-FL compared to non-tFL. **d,** Defining genome-instability strata for paired samples. Scatter plot of average proportion of non-diploid genomic bins with the threshold used to stratify samples into low and high genome instability groups. **e,** Cell-type composition differences by high and low genome instability tFL-DLBCL samples.

## Notes

### Competing Interest Statement

The authors have declared no competing interest.

